# Long-lasting Analgesia via Targeted *in vivo* Epigenetic Repression of Nav1.7

**DOI:** 10.1101/711812

**Authors:** Ana M. Moreno, Glaucilene F. Catroli, Fernando Alemán, Andrew Pla, Sarah A. Woller, Michael Hu, Tony Yaksh, Prashant Mali

## Abstract

Current treatments for chronic pain rely largely on opioids despite their unwanted side effects and risk of addiction. Genetic studies have identified in humans key targets pivotal to nociceptive processing, with the voltage-gated sodium channel, Na_V_1.7 (SCN9A), being perhaps the most promising candidate for analgesic drug development. Specifically, a hereditary loss-of-function mutation in Na_V_1.7 leads to insensitivity to pain without other neurodevelopmental alterations. However, the high sequence similarity between Na_V_ subtypes has frustrated efforts to develop selective inhibitors. Here, we investigated targeted epigenetic repression of Na_V_1.7 via genome engineering approaches based on clustered regularly interspaced short palindromic repeats (CRISPR)-dCas9 and zinc finger proteins as a potential treatment for chronic pain. Towards this end, we first optimized the efficiency of Na_V_1.7 repression *in vitro* in Neuro2A cells, and then by the lumbar intrathecal route delivered both genome-engineering platforms via adeno-associated viruses (AAVs) to assess their effects in three mouse models of pain: carrageenan-induced inflammatory pain, paclitaxel-induced neuropathic pain and BzATP-induced pain. Our results demonstrate: one, effective repression of Na_V_1.7 in lumbar dorsal root ganglia; two, reduced thermal hyperalgesia in the inflammatory state; three, decreased tactile allodynia in the neuropathic state; and four, no changes in normal motor function. We anticipate this genomically scarless and non-addictive **pain** amelioration approach enabling **L**ong-lasting **A**nalgesia via **T**argeted *in vivo* **E**pigenetic **R**epression of Nav1.7, a methodology we dub **pain LATER**, will have significant therapeutic potential, such as for preemptive administration in anticipation of a pain stimulus (pre-operatively), or during an established chronic pain state.

**One sentence summary:** *In situ* epigenome engineering approach for genomically scarless, durable, and non-addictive management of pain.

## INTRODUCTION

Chronic pain affects between 19% to 50% of the world population^1–4^, with more than 100 million people affected in the U.S. alone^5^. Moreover, the number of people reporting chronic pain is expected to increase by 2035 due to the aging global population and prevalence of chronic diseases^6,7^. While chronic pain is more prevalent than cancer, diabetes and cardiovascular disease combined^8^, drug development has not undergone the remarkable progress seen in these other therapeutic areas. Furthermore, current standard of care for chronic pain often relies on opioids, which can have adverse side effects and significant addiction risk. Despite decades of research, the goal of achieving broadly effective, long-lasting, non-addictive therapeutics for chronic pain has remained elusive.

Pain arising from somatic or nerve injury/pathologies typically arises by activation of populations of primary afferent neurons which are characterized by activation thresholds associated with tissue injury and by sensitivity to products released by local tissue injury and inflammation. These afferents terminate in the spinal dorsal horn, where this input is encoded and transmitted by long ascending tracts to the brain, where it is processed into the pain experience. The cell body of a primary afferent lies in its dorsal root ganglion (DRG). These neuronal cell bodies, synthesize the voltage gated sodium channels that serve to initiate and propagate the action potential^9–11^. While local anesthetics can yield a dense anesthesia, previous work has in fact shown that nonspecific sodium channel blockers such as lidocaine delivered systemically at subanesthetic concentrations were able to have selective effects upon hyperpathia in animal models and humans^12–15^.

It is now known that there are nine voltage-gated sodium channel subtypes along with numerous splice variants. Of note, three of these isotypes: Na_V_1.7^16,17^, Na_V_1.8^18–20^, and Na_V_1.9^21,22^ have been found to be principally expressed in primary afferent nociceptors. The relevance of these isotypes to human pain has been suggested by the observation that a loss-of-function mutation in Na_V_1.7 (SCN9A) leads to congenital insensitivity to pain (CIP), a rare genetic disorder. Conversely, gain of function mutations yield anomalous hyperpathic states^23–29^. Based on these observations, the Na_V_1.7 channel has been considered an attractive target for addressing pathologic pain states and for developing chronic pain therapies^17,29–31^. Efforts to develop selective small molecule inhibitors have, however, been hampered due to the sequence similarity between Na_V_ subtypes. Many small-molecule drugs targeting Na_V_1.7 have accordingly failed due to side effects caused by lack of targeting specificity or their bioavailability by the systemic route^32^. Additionally, antibodies have faced a similar situation, since there is a tradeoff between selectivity and potency due to the binding of a specific (open or close) conformation of the channel, with binding not always translating into successful channel inhibition^33–36^. Further, it is not clear that such antibodies can gain access to the appropriate Na_V_1.7 channels and yield a reliable block of their function. Consequently, in spite of preclinical studies demonstrating that decreased Na_V_1.7 activity leads to a reduction in inflammatory and neuropathic pain^16–22,37^, no molecule targeting this gene product has reached the final phase of clinical trials^32^. We therefore took an alternative approach by epigenetically modulating the expression of Na_V_1.7 using two genome engineering tools, clustered regularly interspaced short palindromic repeats (CRISPR)– Cas9 (CRISPR-Cas9) and zinc-finger proteins (ZFP), such that one could engineer highly specific, long-lasting and reversible treatments for pain.

Through its ability to precisely target disease-causing DNA mutations, the CRISPR-Cas9 system has emerged as a potent tool for genome manipulation, and has shown therapeutic efficacy in multiple animal models of human diseases^38–44^. However, permanent genome editing, leading to permanent alteration of pain perception, may not be desirable. For example, pain can be a discomforting sensory and emotional experience, but it plays a critical role alerting of tissue damage. Permanent ablation could thus have detrimental consequences. For these reasons, we have employed a catalytically inactivated “dead” Cas9 (dCas9, also known as CRISPRi), which does not cleave DNA but maintains its ability to bind to the genome via a guide-RNA (gRNA), and fused it to a repressor domain (Krüppel-associated box, KRAB) to enable non-permanent gene repression of Na_V_1.7. Previously, we and others have shown that through addition of a KRAB epigenetic repressor motif to dCas9, gene repression can be enhanced with a high level of specificity both *in vitro*^45–49^ and *in vivo*^50,51^. This transcriptional modulation system takes advantage of the high specificity of CRISPR-Cas9 while simultaneously increasing the safety profile, as no permanent modification of the genome is performed. As a second approach for *in situ* epigenome repression of Na_V_1.7, we also utilized zinc-finger-KRAB proteins (ZFP-KRAB), consisting of a DNA-binding domain made up of Cys_2_His_2_ zinc fingers fused to a KRAB repressor. ZFP constitute the largest individual family of transcriptional modulators encoded by the genomes of higher organisms^52^, and with prevalent synthetic versions engineered on human protein chasses present a potentially low immunogenicity *in vivo* targeting approach^53–55^. We sought to produce a specific anatomic targeting of the gene regulation by delivering both epigenetic tools in an adeno-associated virus (AAV) construct into the spinal intrathecal space. Of note, many AAVs have been shown to produce a robust transduction of the dorsal root ganglion^56–58^. This approach has several advantages as it permits the use of minimal viral loads and reduces the possibility of systemic immunogenicity.

Since pain perception is etiologically diverse and multifactorial, several rodent pain models have been utilized to study pain signaling and pain behaviors^59^. In the present work we sought to characterize the effects of CRISPR-mediated knock down of Na_V_1.7 using three mechanistically distinct models: i) thermal sensitivity in control (normal) and unilateral inflammation-sensitized hind paw; ii) a poly neuropathy induced by a chemotherapeutic yielding a bilateral hind paw tactile allodynia and, iii) a spinally evoked bilateral hind paw tactile allodynia induced by spinal activation of purine receptors. Pain due to tissue injury and inflammation results from a release of factors that sensitize the peripheral terminal of the nociceptive afferent neuron. This phenotype can be studied through local application of carrageenan to the paw resulting in inflammation, swelling, increased expression of Na_V_1.7^60^ and a robust increase in thermal and mechanical sensitivity (hyperalgesia)^61,62^. Chemotherapy to treat cancer often leads to a polyneuropathy characterized by increased sensitivity to light touch (e.g. tactile allodynia) and cold. Paclitaxel is a commonly used chemotherapeutic that increases the expression of Na_V_ 1.7 in the nociceptive afferents^63,64^ and induces a robust allodynia in the animal models^65–67^. Finally, ATP (adenosine triphosphate) by an action on a variety of purine receptors expressed on afferent terminals and second order neurons and non neuronal cells has been broadly implicated in inflammatory, visceral and neuropathic pain states^57–60^. Thus, intrathecal delivery of a stable ATP analogue (BzATP: 2’,3’-O-(4-benzoylbenzoyl)-ATP) results in a long-lasting allodynia in mice^71,72^.

Thus, we first tested various KRAB-dCas9 and ZFP-KRAB constructs in a mouse neuroblastoma cell line that expresses Na_V_1.7 (Neuro2a) and optimized repression levels. We next packaged the constructs with the strongest *in vitro* repression into AAV9 and injected these intrathecally into adult C57BL/6J mice. After 21 days, we induced paw inflammation via injection of carrageenan, and tested for thermal hyperalgesia. Our results demonstrated *in vivo* repression of Na_V_1.7 and a decrease in thermal hyperalgesia. Similarly, we tested our epigenome strategy in two neuropathic pain models: chemotherapy-induced (paclitaxel) and BzATP-induced neuropathic pain. The results in the paclitaxel-induced neuropathic pain model indicate that repression of Na_V_1.7 leads to reduced tactile and cold allodynia. In addition, KRAB-dCas9 injected mice showed reduced tactile allodynia after administration of the ATP analogue BzATP. As many pain states occurring after chronic inflammation and nerve injury represent an enduring condition, typically requiring constant re-medication, these genetic approaches provide ongoing and controllable regulation of this aberrant processing. Overall, these *in situ* epigenetic approaches could represent a viable replacement for opioids and serve as a potential therapeutic approach for long lasting chronic pain.

## RESULTS

### In vitro optimization of epigenetic engineering tools to enable Na_V_1.7 repression

With the goal of developing a therapeutic product that relieves chronic pain in a non-permanent, non-addictive and long-lasting manner, we explored the use of two independent genetic approaches to inhibit the transmission of pain at the spinal level **(Figure 1a)**. To establish robust Na_V_1.7 repression, we first compared *in vitro* repression efficacy of Na_V_1.7 using KRAB-dCas9 and ZFP-KRAB constructs. Towards this, we cloned ten guide-RNAs (gRNAs; **Supplementary Table 1**) —designed by an *in silico* tool^73^ that predicts highly effective gRNAs based on chromatin position and sequence features— into our previously developed split-dCas9 platform^50^. We also cloned the two gRNAs that were predicted to have the highest efficiency (SCN9A-1 and SCN9A-2) into a single construct, since we have previously shown that higher efficacy can be achieved by using multiple gRNAs^50^. We next utilized four ZFP-KRAB constructs targeting the Na_V_1.7 DNA sequence **(Supplementary Table 2)**. These constructs were transfected into a mouse neuroblastoma cell line that expresses Na_V_1.7 (Neuro2a) and we confirmed repression of Na_V_1.7 relative to GAPDH with qPCR. Six of ten gRNAs repressed the Na_V_1.7 transcript by >50% compared to the non-targeting gRNA control, with gRNA-2 being the single gRNA having the highest repression (56%) and with the dual-gRNA having repression levels of 71% (p < 0.0001), which we utilized for subsequent *in vivo* studies **(Supplementary Figure 1a)**. Of the ZFP-KRAB designs, the Zinc-Finger-4-KRAB construct had the highest repression (88%; p < 0.0001) compared to the negative control (mCherry), which we chose for subsequent *in vivo* studies **(Supplementary Figure 1a)**. Western blotting confirmed a corresponding decrease in protein level for both the Zinc-Finger-4-KRAB and KRAB-dCas9-dual-gRNA groups **(Supplementary Figure 1b)**.

**Figure 1:**
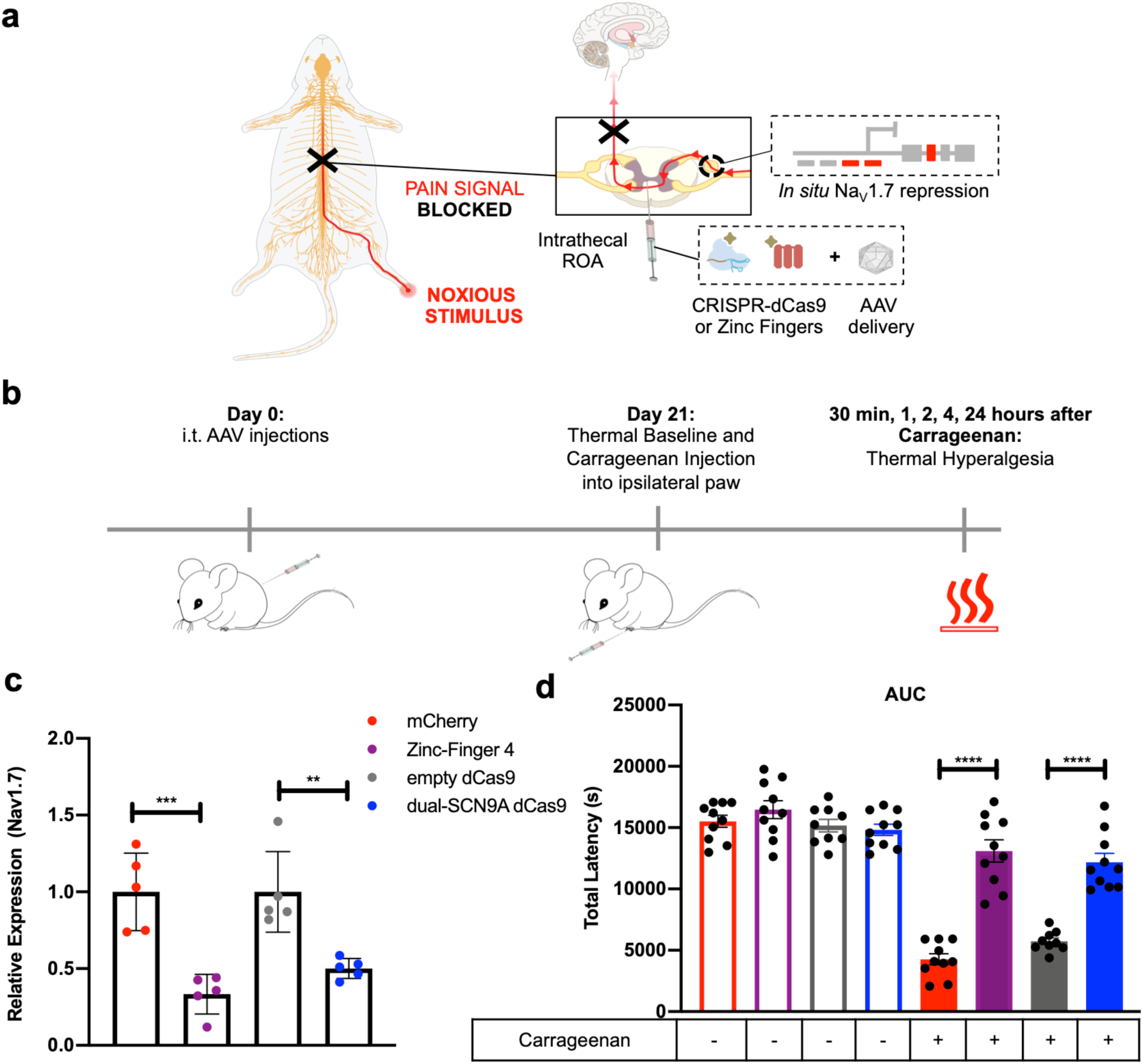
*In situ* repression of Na_V_1.7 leads to pain amelioration in a carrageenan model of inflammatory pain. **(a)** Schematic of the overall strategy used for *in situ* Na_V_1.7 repression using ZFP-KRAB and KRAB-dCas9. Na_V_1.7 is a DRG channel involved in the transduction of noxious stimuli into electric impulses at the peripheral terminals of DRG neurons. *In situ* repression of Na_V_1.7 via AAV-ZFP-KRAB and AAV-KRAB-dCas9 is achieved through intrathecal injection leading to disruption of the pain signal before reaching the brain. **(b)** Schematic of the carrageenan-induced inflammatory pain model. At day 0, mice were intrathecally injected with either AAV9-Zinc-Finger-4-KRAB, AAV9-mCherry, AAV9-KRAB-dCas9-dual-gRNA or AAV9-KRAB-dCas9-no-gRNA. 21 days later, thermal pain sensitivity was measured in all mice with the Hargreaves assay. In order to establish a baseline level of sensitivity, mice were tested for tactile threshold using von Frey filaments before carrageenan injection. Mice were then injected with carrageenan in their left paw (ipsilateral) while the right paw (contralateral) was injected with saline as an in-mouse control. They were then tested for thermal paw-withdrawal latency at 30 min, 1, 2, 4, and 24 hours after carrageenan administration. **(c)** *In vivo* Na_V_1.7 repression efficiencies: Twenty-four hours after carrageenan administration, mice DRG (L4-L6) were harvested and Na_V_1.7 repression efficacy was determined by qPCR. (n=5; error bars are SEM; Student’s t-test; ***p = 0.0008, **p=0.0033). **(d)** The aggregate paw withdrawal latency was calculated as area under the curve (AUC) for both carrageenan and saline injected paws. Mice treated with Zinc-Finger-4-KRAB and KRAB-dCas9-dual-gRNA had significant increased paw-withdrawal latencies in carrageenan-injected paws (n=10; error bars are SEM; Student’s t-test, ****p < 0.0001).

### In vivo evaluation of ZFP-KRAB and KRAB-dCas9 treatment efficacy in a carrageenan model of inflammatory pain

Having established *in vitro* Na_V_1.7 repression, we next focused on testing the effectiveness of the best ZFP-KRAB and KRAB-dCas9 constructs from the *in vitro* screens (Zinc-Finger-4-KRAB and KRAB-dCas9-dual-gRNA) in a carrageenan-induced model of inflammatory pain. Mice were intrathecally (i.t.) injected with 1E+12 vg/mouse of AAV9-mCherry (negative control; n=10), AAV9-Zinc-Finger-4-KRAB (n=10), AAV9-KRAB-dCas9-no-gRNA (negative control; n=10) and AAV9-KRAB-dCas9-dual-gRNA (n=10). The intrathecal delivery of AAV9, which has significant neuronal tropism^58^, serves to efficiently target DRG (**Supplementary Figure 2a)**. After 21 days, thermal pain sensitivity was measured to establish a baseline response threshold. Inflammation was induced in all four groups of mice by injecting one hind paw with carrageenan (ipsilateral), while the other hind paw (contralateral) was injected with saline to serve as an in-mouse control. Mice were then tested for thermal pain sensitivity at 30 minutes, 1, 2, 4, and 24 hours after carrageenan injection **(Figure 1b)**. Twenty-four hours after carrageenan administration, mice were euthanized and DRG (L4-L6) were extracted. The expression levels of Na_V_1.7 was determined by qPCR, and a significant repression of Na_V_1.7 was observed in mice injected with AAV9-Zinc-Finger-4-KRAB (67%; p=0.0008) compared to mice injected with AAV9-mCherry, and in mice injected with AAV9-KRAB-dCas9-dual-gRNA (50%; p=0.0033) compared to mice injected with AAV9-KRAB-dCas9-no-gRNA **(Figure 1c)**. The mean paw withdrawal latencies (PWL) were calculated for both carrageenan and saline injected paws **(Supplementary Figure 2b, c)** and the area under the curve (AUC) for the total mean PWL was calculated. As expected, compared to saline-injected paws, carrageenan-injected paws developed thermal hyperalgesia, measured by a decrease in PWL after application of a thermal stimulus **(Figure 1d)**. We also observed a significant increase in PWL in mice injected with either AAV9-Zinc-Finger-4-KRAB or AAV9-KRAB-dCas9-dual-gRNA, indicating that the repression of Na_V_1.7 in mouse DRG leads to lower thermal hyperalgesia in an inflammatory pain state. The thermal latency of the control (un-inflamed paw) was not significantly different across AAV treatment groups, indicating that the knock down of the Na_V_1.7 had minimal effect upon normal thermal sensitivity. As an index of edema/inflammation, we measured the ipsilateral and contralateral paws with a caliper before and 4 hours after carrageenan injection, which is the time point with the highest thermal hyperalgesia. We observed significant edema formation in both experimental and control groups, indicating that Na_V_1.7 repression has no effect on inflammation **(Supplementary Figure 2d).**

### Benchmarking in vivo ZFP-KRAB treatment efficacy with established small molecule drug gabapentin in a carrageenan model of inflammatory pain

To validate the efficacy of ZFP-KRAB in ameliorating thermal hyperalgesia in a carrageenan model of inflammatory pain, we next conducted a separate experiment and tested the small molecule drug gabapentin as a positive control. Mice were i.t. injected with 1E+12 vg/mouse of AAV9-mCherry (n=5), AAV9-Zinc-Finger-4-KRAB (n=6), or saline (n=5). After 21 days, thermal nociception was measured in all mice as previously described. One hour before carrageenan administration, the mice that received intrathecal saline were injected as a positive comparator with intraperitoneal (i.p.) gabapentin (100 mg/kg). This agent is known to reduce carrageenan-induced thermal hyperalgesia in rodents through binding to spinal alpha2 delta subunit of the voltage gated calcium channel^74,75^. Twenty-four hours after carrageenan administration, mice were euthanized and DRG (L4-L6) were extracted. The expression levels of Na_V_1.7 were determined by qPCR, and a significant repression of Na_V_1.7 was observed in AAV9-Zinc-Finger-4-KRAB (***p=0.0007) and in the gabapentin groups (*p=0.0121) **(Supplementary Figure 3a)**. We measured the ipsilateral and contralateral paws with a caliper before and 4 hours after carrageenan injection, and confirmed significant edema formation in the injected paw of all groups as compared to the non-injected paw in all groups **(Supplementary Figure 3b)**. The mean PWL was calculated for both carrageenan and saline injected paws **(Figure 2b, c)**. We then compared paw withdrawal latencies of carrageenan injected paws for AAV9-Zinc-Finger-4-KRAB and gabapentin groups at each time point to the AAV9-mCherry carrageenan injected control using a two-way ANOVA calculation to determine whether there was any significant reduction in thermal hyperalgesia **(Supplementary Figure 3c)**. When comparing carrageenan-injected hind paws, we observed that only AAV9-Zinc-Finger-4-KRAB had significantly higher PWL at all the time points following carrageenan injection when compared to the AAV9-mCherry control. We also observed significance in PWL for the gabapentin positive control group at the 30 minute, 1 hour, and four hour time points, but not the 24 hour time point. This result reflects the half-life of gabapentin (3-5 hours). We then calculated the area under the curve (AUC) for thermal hyperalgesia. We observed a significant increase in PWL in the carrageenan-injected gabapentin group (p=0.0208) (**Figure 2b**), and in the Zinc-Finger-4-KRAB group (115% improvement, p=0.0021) **(Figure 2c)** compared to the carrageenan-injected AAV9-mCherry control. In addition, the AAV9-Zinc-Finger-4-KRAB group had 31% higher PWL than the gabapentin positive control group. Of note, the thermal escape latency of the contralateral non-inflamed paw showed no significant difference among groups.

**Figure 2:**
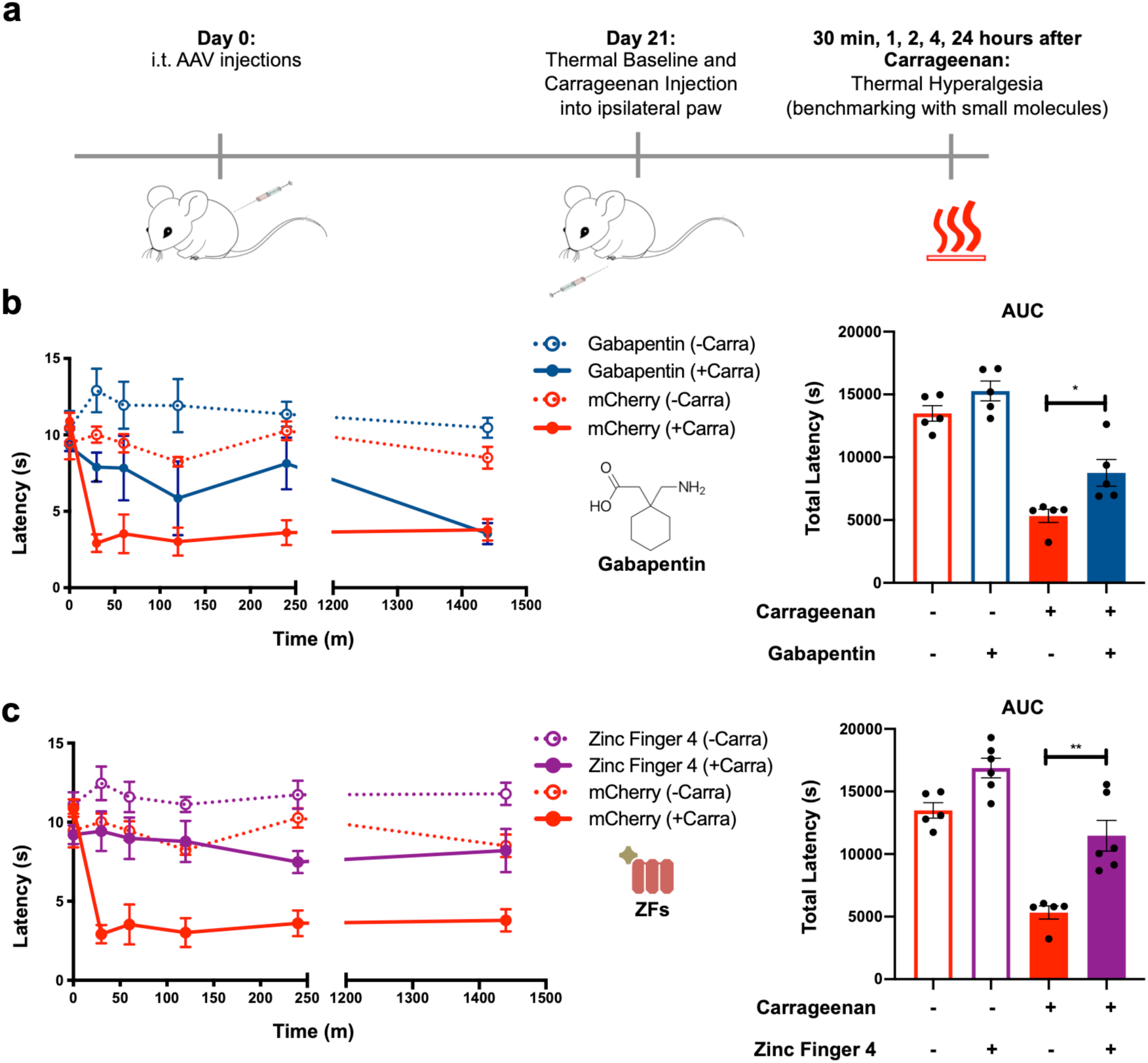
Benchmarking of *in situ* repression of Na_V_1.7 using Zinc-Finger-KRAB with established small molecule drug gabapentin. **(a)** Schematic of the experimental approach. **(b)** Time course of thermal hyperalgesia after the injection of carrageenan (solid lines) or saline (dotted lines) into the hind paw of mice injected with gabapentin (100mg/kg) and mCherry and Zinc-Finger-4-KRAB are plotted. Mean paw withdrawal latencies (PWL) are shown. The AUC of the thermal-hyperalgesia time-course are plotted on the right panels. A significant increase in PWL is seen in the carrageenan-injected paws of mice injected with gabapentin and Zinc-Finger-4-KRAB (n=5 for mCherry and gabapentin and n=6 for Zinc-Finger-4-KRAB; error bars are SEM; Student’s t-test, *p = 0.0208, **p=0.0021).

### In vivo repression of Na_V_1.7 leads to amelioration of pain in a poly neuropathic pain model

After having established *in vivo* efficacy in an inflammatory pain model, we went on to validate our epigenome repression strategy for neuropathic pain using the polyneuropathy model by the chemotherapeutic paclitaxel. To establish this model, mice were first injected with 1E+12 vg/mouse of AAV9-mCherry (n=8), AAV9-Zinc-Finger-4-KRAB (n=8), AAV9-KRAB-dCas9-dual-gRNA (n=8), AAV9-KRAB-dCas9-no-gRNA (n=8), or saline (n=16). 14 days later and before paclitaxel administration, we established a baseline for tactile threshold (von Frey filaments). Mice were then administered paclitaxel at days 14, 16, 18, and 20, with a dosage of 8 mg/kg (total cumulative dosage of 32 mg/kg), with a group of saline injected mice not receiving any paclitaxel (n=8) to establish the tactile allodynia caused by the chemotherapeutic. 21 days after the initial injections and one hour before testing, a group of saline injected mice (n=8) were injected with i.p. gabapentin (100 mg/kg). Mice were then tested for tactile allodynia via von Frey filaments and for cold allodynia via acetone testing **(Figure 3a)**. A 50% tactile threshold was calculated. We observed a decrease in tactile threshold in mice receiving AAV9-mCherry and AAV9-KRAB-dCas9-no-gRNA, while mice that received gabapentin, AAV9-Zinc-Finger-4-KRAB, and AAV9-KRAB-dCas9-dual-gRNA had increased withdrawal thresholds, indicating that *in situ* Na_V_1.7 repression leads to amelioration in chemotherapy induced tactile allodynia **(Figure 3b)**. Similarly, an increase in the number of withdrawal responses is seen in mice tested for cold allodynia in the negative control groups (AAV9-mCherry and AAV9-KRAB-dCas9-no-gRNA), while both AAV9-Zinc-Finger-4-KRAB and AAV9-KRAB-dCas9-dual-gRNA groups had a decrease in withdrawal responses, indicating that *in situ* repression of Na_V_1.7 also leads to a decrease in chemotherapy induced cold allodynia **(Figure 3c)**.

**Figure 3:**
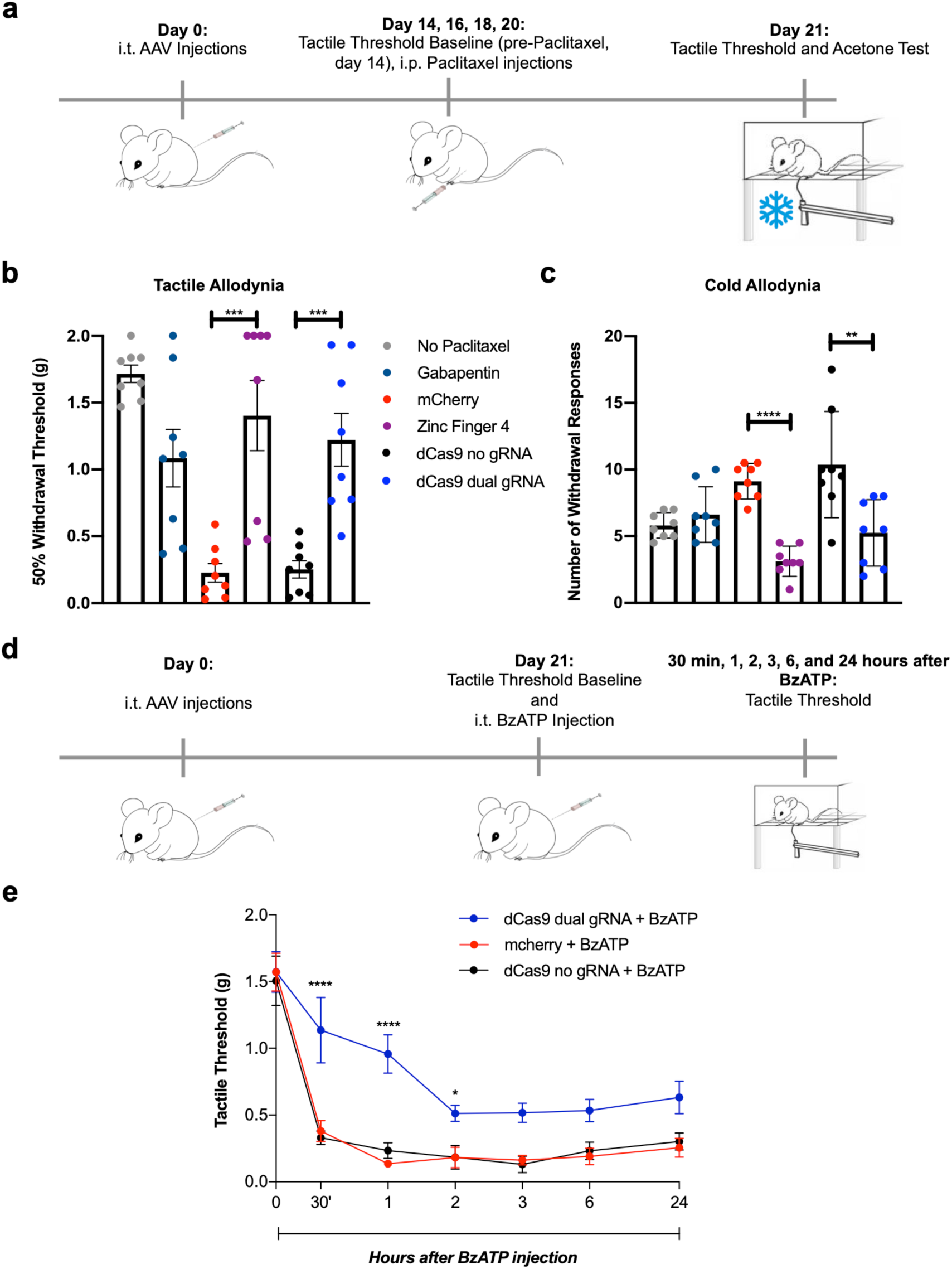
*In vivo* efficacy of Zinc-Finger-KRAB and KRAB-dCas9 in two neuropathic pain models. **(a)** Schematic of the paclitaxel-induced neuropathic pain model. Mice were i.t. injected with AAV9-mCherry, AAV9-Zinc-Finger-4-KRAB, AAV9-KRAB-dCas9-no-gRNA, AAV9-KRAB-dCas9-dual-gRNA or saline. Following baseline von Frey threshold testing at day 14, mice were then injected i.p. with 8mg/kg of paclitaxel at 14, 16, 18, 20 days after i.t. injection. 21 days after i.t. injection, mice were tested for tactile allodynia via von Frey filaments and for cold allodynia via the application of acetone. **(b)** *In situ* repression of Na_V_1.7 via Zinc-Finger-4-KRAB and KRAB-dCas9-dual-gRNA reduces paclitaxel-induced tactile allodynia. (n=8; error bars are SEM; Student’s t-test; ***p = 0.0007, ***p = 0.0004) **(c)** *In situ* repression of Na_V_1.7 via Zinc-Finger-4-KRAB and KRAB-dCas9-dual-gRNA reduces paclitaxel-induced cold allodynia. (n=8; error bars are SEM; Student’s t-test; ****p < 0.0001, **p = 0.008). **(d)** Schematic of the BzATP pain model. Mice were injected at day 0 with AAV9-mCherry, AAV9-KRAB-dCas9-no-gRNA or AAV9-KRAB-dCas9-dual-gRNA. 21 days later, mice were baselined for von Frey tactile threshold and were then i.t. injected with 30 nmol BzATP. 30 minutes, 1, 2, 3, 6, and 24 hours after BzATP administration, mice were tested for tactile allodynia. **(e)** *In situ* repression of Nav1.7 via KRAB-dCas9-dual-gRNA reduces tactile allodynia in a BzATP model of neuropathic pain (n=5 for KRAB-dCas9-no-gRNA and n=6 for the other groups, two-way ANOVA with Bonferonni *post-hoc* test; ****p < 0.0001, *p = 0.0469).

### In vivo repression of Na_V_1.7 leads to amelioration of pain in a model of spinal evoked pain

We next tested whether *in situ* repression of Na_V_1.7 via KRAB-dCas9 could ameliorate neuropathic pain induced by BzATP. This molecule activates P2X receptors located on central terminals leading to a centrally mediated hyperalgesic state. We first injected mice with 1E+12 vg/mouse of AAV9-mCherry (n=6), AAV9-KRAB-dCas9-no-gRNA (n=5), and AAV9-KRAB-dCas9-dual-gRNA (n=6). After 21 days, tactile thresholds were determined with von Frey filaments, and mice were injected i.t. with BzATP (30 nmol). Tactile allodynia was then measured at 30 min, 1, 2, 3, 6, and 24 hours after BzATP administration **(Figure 3d)**. We observed a significant decrease in tactile allodynia at 30 min, 1 and 2 hour time points in mice injected with AAV9-KRAB-dCas9-dual-gRNA, and an overall increase in tactile threshold at all time points **(Figure 3e)**.

### Durable pain amelioration via the in situ repression of Na_V_1.7 with Zinc-Finger-KRAB and KRAB-dCas9

To determine whether *in situ* repression of Na_V_1.7 was efficacious long-term, we repeated the carrageenan inflammatory pain model and tested thermal hyperalgesia at 21 and 42 days after i.t. AAV injection (n=8/group) **(Figure 4a)**. We observed a significant improvement in PWL for carrageenan-injected paws in Zinc-Finger-4-KRAB groups at both day 21 **(Supplementary Figure 3d)** and day 42 **(Figure 4b)** demonstrating the durability of this approach. To determine whether *in situ* repression of Na_V_1.7 was also efficacious long-term in a poly neuropathic pain model, we measured tactile and cold allodynia 49 days after initial AAV injections and 29 days after the last paclitaxel injection (total cumulative dosage of 32 mg/kg; **Figure 4c**). Compared to the earlier time point **(Figure 3b, c)**, we observed that mice from both AAV9-mCherry (n=8) and AAV9-KRAB-dCas9-dual-gRNA (n=8) groups had increased tactile allodynia at day 49 as compared to day 21, and responded to the lowest von Frey filament examined (0.04 g). In comparison, mice receiving AAV9-Zinc-Finger-4-KRAB and AAV9-KRAB-dCas9-dual-gRNA had increased withdrawal thresholds, indicating that *in situ* Na_V_1.7 repression leads to long-term amelioration in chemotherapy-induced tactile allodynia **(Figure 4d)**. As before, an increase in the number of withdrawal responses is seen in mice tested for cold allodynia in the negative control groups (AAV9-mCherry and AAV9-KRAB-dCas9-no-gRNA), while both AAV9-Zinc-Finger-4-KRAB and AAV9-KRAB-dCas9-dual-gRNA groups had a decrease in withdrawal responses, indicating that *in situ* repression of Na_V_1.7 also leads long-term amelioration of chemotherapy induced cold allodynia **(Figure 4e)**

**Figure 4:**
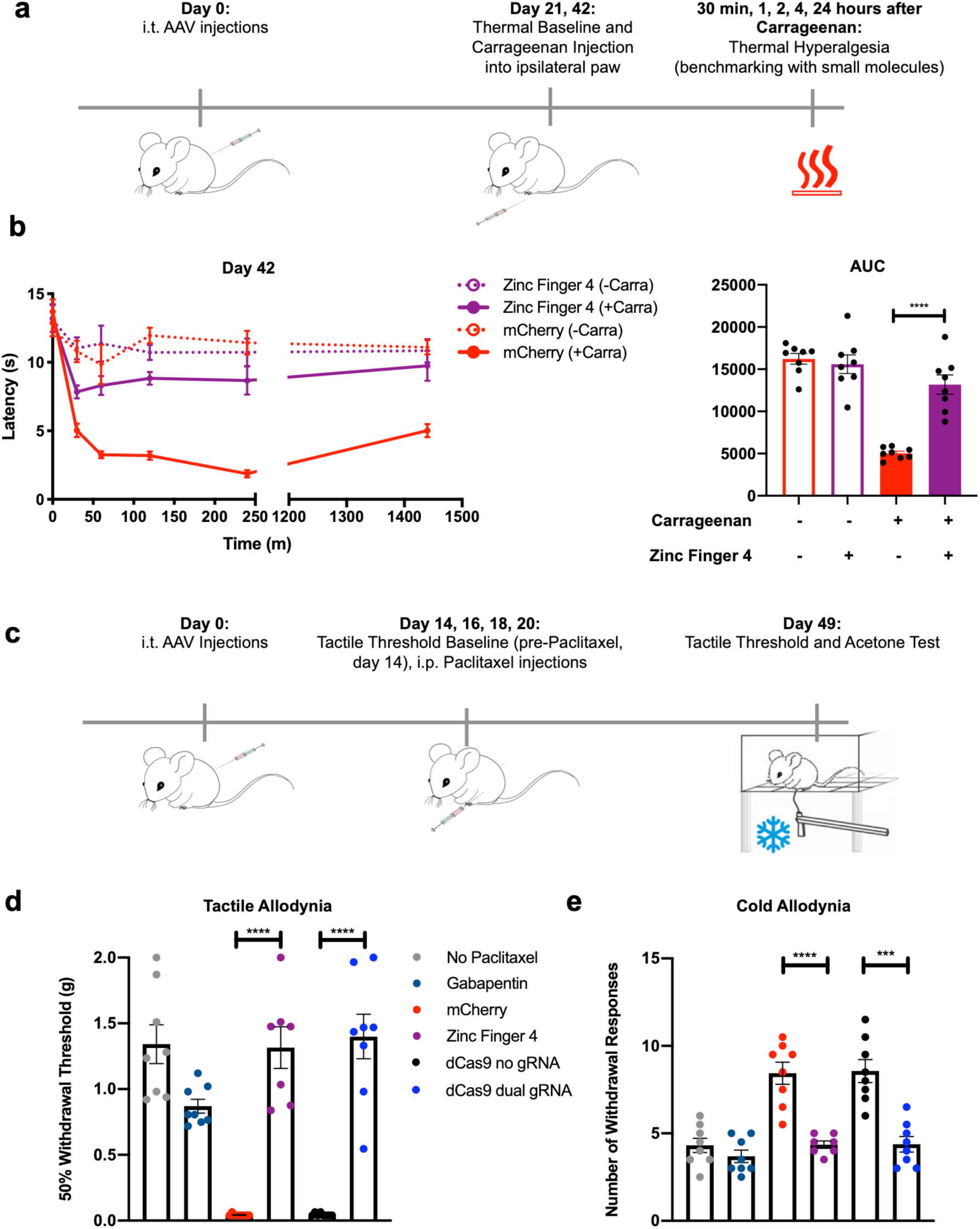
Long-term efficacy of Zinc-Finger-KRAB and KRAB-dCas9 in two independent pain models. **(a)** Timeline of the carrageenan-induced inflammatory pain model. **(b)** Time course of thermal hyperalgesia after the injection of carrageenan (solid lines) or saline (dotted lines) into the hind paw of mice 42 days after i.t. injection with AAV9-mCherry and AAV9-Zinc-Finger-4-KRAB are plotted. Mean paw withdrawal latencies are shown. The AUC of the thermal-hyperalgesia time-course are plotted on the right panel. A significant increase in PWL is seen in the carrageenan-injected paws of mice injected with AAV9-Zinc-Finger-4-KRAB (n=8; error bars are SEM; Student’s t-test; ****p < 0.0001). **(c)** Schematic of the paclitaxel-induced neuropathic pain model. **(d)** *In situ* repression of Na_V_1.7 via Zinc-Finger-4-KRAB and KRAB-dCas9-dual-gRNA reduces paclitaxel-induced tactile allodynia 49 days after last paclitaxel injection (n=7 for Zinc-Finger-4-KRAB and n=8 for other groups; error bars are SEM; Student’s t-test; ****p < 0.0001). **(e)** *In situ* repression of Na_V_1.7 via Zinc-Finger-4-KRAB and KRAB-dCas9-dual-gRNA reduces paclitaxel-induced cold allodynia. (n=7 for Zinc-Finger-4-KRAB and n=8 for other groups; error bars are SEM; Student’s t-test; ****p < 0.0001, ***p = 0.0001).

## DISCUSSION

In this study, we investigated the efficacy of the repression of Na_V_1.7 in the dorsal root ganglia using two independent genome engineering platforms —KRAB-dCas9 and Zinc-Finger-KRAB proteins— to treat acute and persistent nociceptive processing generated in murine models of peripheral inflammation and poly neuropathy. We believe the promising results suggest the utility of the approach for developing a therapeutic reagent.

### Development of targeted constructs

The work employed multiple guide-RNAs (gRNAs) clones which were rationally designed using an *in silico* tool^73^ that predicts effective gRNAs based on chromatin position and sequence features into the split-dCas9 platform^50^. Similarly, we devised multiple ZFP-KRAB Na_V_1.7 DNA targeting constructs. These constructs were transfected into a murine neuroblastoma cell line that expresses Na_V_1.7 (Neuro2a) and repression of Na_V_1.7 was confirmed. Constructs showing the highest level of repression were chosen for subsequent *in vivo* studies.

Although other technologies, such as RNAi have been utilized to target Na_V_1.7, a recent study has shown that the off-target effects of RNAi, as compared to CRISPRi, are far stronger and more pervasive than generally appreciated^76,77^. In addition, as an exogenous system, CRISPR and ZFPs (unlike RNAi) do not compete with endogenous machinery such as microRNA or RISC complex function. Thus, RNAi can have an impact in the regular homeostatic mechanisms of RNA synthesis and degradation. In addition, CRISPR and ZFP methods target genomic DNA instead of RNA, which means that to achieve an effect, RNAi methods require a higher dosage with poorer pharmacokinetics prospects, as there is usually a high RNA turnover^46^.

### In vivo spinal Na_V_1.7 knock down

In these studies, we found that mice injected with either epigenetic platform had significantly reduced DRG expression of Na_V_1.7. Other studies have shown that partial repression of Na_V_1.7^78–83^ is sufficient to ameliorate pain. This knock down serves to produce a significant reversal of the hyperalgesia induced by hind paw inflammation. Using antisense oligonucleotides, mechanical pain could be ameliorated with 30 to 80% Na_V_1.7 repression levels^81^. Using microRNA 30b, around 50% repression of Na_V_1.7 relieved neuropathic pain^82^, while more recently microRNA182 ameliorated pain preventing Na_V_1.7 overexpression in spared nerve injury rats^79^. Similarly, shRNA mediated knockdown of Na_V_1.7 prevented its overexpression in burn injury relieving pain^78^. Other studies did not quantify the Na_V_1.7 repression levels needed to reduce pain^80^. Additionally, shRNA lentiviral vectors can reduce bone cancer pain by repressing Na_V_1.7 40 to 60%^83^. Further studies are needed to determine what the minimum dosage to have an effect is.

The role of Na_V_1.7 has been implicated in a variety of preclinical models, including those associated with robust inflammation as in the rodent carrageenan and CFA model. We examined the effect of knocking down Na_V_1.7 in a paclitaxel-induced poly neuropathy. Previous work has shown that this treatment will induce Na_V_1.7^64,87^. Both epigenetic repressors ameliorate tactile allodynia to the same extent as the internal comparator gabapentin. Finally, we further addressed the role of Na_V_1.7 knock down in hyperpathia induced by i.t. injection of BzATP. This was significantly attenuated in mice previously treated with KRAB-dCas9. Spinal purine receptors have been shown to play a pivotal role in the nociceptive processing initiated by a variety of stimulus conditions including inflammatory/incisional pain and a variety of neuropathies^72,88–90^. The present observations suggest that the repression of afferent Na_V_1.7 expression in the nociceptor leads to a suppression of enhanced tactile sensitivity induced centrally. The mechanism underlying these results may reflect upon the observation that down regulation of Na_V_1.7 in the afferent may serve to minimize the activation of microglia and astrocytes^83^. These results suggest that, at least partially, pain signal transduction through Na_V_1.7 is downstream of ATP signaling.

We chose gabapentin as a positive control due to evidence that it decreases carrageenan-induced thermal hyperalgesia in rodents and because it is known to repress Na_V_1.7. Our results are consistent with previous studies that showed a inhibitory effect of gabapentin on Na_V_1.7 expression levels, ultimately leading to a reduction of neuronal excitability^91^. In addition, although it has been previously reported that gabapentin can lower inflammation on rat paw edema induced by carrageenan^92^, we did not find any reduction in inflammation in the gabapentin group This could be due to a difference in animal model, gabapentin dosage, or the concentration of carrageenan injected. Because only one dosage of gabapentin was utilized for these experiments, an additional group of mice receiving a different gabapentin dosage or with a second dosage can be utilized as a secondary positive control.

Of note, these drug effects examined in the polyneuropathy and carrageenan model appeared to persist unchanged for at least 6-7 weeks. Long term expression has been similarly noted in other gene therapy studies^57,93–95^. Importantly, these effects were unaccompanied by any detectable adverse motor effects or changes in bladder function.

### Spinal route for therapeutic delivery

The presented research shows efficacy of spinal reduction in Na_V_1.7 in three models of hyperpathia using two complementary epigenetic tools. These studies clearly established significant target engagement and clear therapeutic efficacy with no evident adverse events after intrathecal knock down of Na_V_1.7 with two independent platforms. The intrathecal route of delivery represents an appropriate choice for this therapeutic approach. The role played by Na_V_1.7 is in the nociceptive afferents, and their cell bodies are in the respective segmental DRG neurons. Accordingly, the DRG represents the target for this transfection motif. The intrathecal delivery route efficiently place AAVs to the DRG neurons which minimizes the possibility of off target biodistribution and reduces the viral load required to get transduction. Importantly, the relative absence of B and T cells^96,97^ in the cerebrospinal fluid, reduces the potential immune response. In this regard, as ZFPs are engineered on human protein chasses, they intrinsically constitute a targeting approach with even lower potential immunogenicity. Indeed, a study in non-human primates (NHP) found that intrathecal delivery of a non-self protein (AAV9-GFP) produced immune responses which were not seen with the delivery of a self-protein^98^.

### Future directions

These results displaying target engagement and efficacy provide strong support for the development of these platforms for pain control. Several issues are pertinent: One, further studies are needed to determine what is the minimum effective AAV dosage to produce knock down and therapeutic effects. Two, this work showed reduction in Na_V_1,7 at 21, 42, and 49 days after AAV injections and corresponding changes in pain behavior. However, still longer-term studies need to be performed to evaluate the actual duration of treatment and whether any compensatory mechanisms take place due to Na_V_1.7 repression. Three, further studies must be performed to explore the properties of repeat-dosing. Four, we validated our approach in three mouse pain models. However, other models of inflammatory pain should be tested to further validate our results. Carrageenan produces a model of persistent pain and hyperalgesia that best represents an acute phase from 1-24 hours and that converts to chronic inflammation by two weeks^61^. Therefore, the performed experiments can be repeated two-weeks after administering carrageenan to determine efficacy in a chronic inflammatory pain state. The Complete Freund’s adjuvant (CFA) model, collagen type II antibodies (CAIA) or K/BxN transgenic mice mimic the pathology of rheumatoid arthritis such as: widespread inflammation with the greatest effect distally, cartilage degradation, and elevated inflammatory cytokines in the joint fluid, and are thus additional important models to explore^59^. Focus on visceral pain will also be an important direction moving forward. Finally, five, other species including rats must be explored to further validate this approach.

### Therapeutic utility of intrathecal CRISPR/ZFP

As a potential clinical treatment, KRAB-dCas9 and ZFP-KRAB show promise for treating chronic inflammatory and neuropathic pain. These systems allow for potentially reversible gene therapy, which is advantageous in the framework of chronic pain, as permanent pain insensitivity is not desired. Additionally, the weeks-long duration presents a significant advantage compared to existing drugs which must be taken daily or hourly, and which may have undesirable addictive effects. Taken together, the results of these studies show a promising new avenue for treatment of chronic pain, a significant and increasingly urgent issue in our society. For instance, cancer treatment related side effects, and in particular chemotherapy induced polyneuropathy is one of the most common adverse events which could potentially be targeted using this system^101,102^. In this instance, a therapeutic approach that endures for months is preferable to one that is irreversible. Furthermore, the use of multiple neuraxial interventions over time is a common motif for clinical interventions as with epidural steroids where repeat epidural delivery may occur over the year at several month intervals^103^. Taken together, we anticipate this genomically scarless and non-addictive pain amelioration approach enabling long-lasting analgesia via targeted *in vivo* epigenetic repression of Nav1.7, will have significant therapeutic potential, such as for preemptive administration in anticipation of a pain stimulus (pre-operatively), or during an established chronic pain state.

## ACKNOWLEDGEMENTS

We thank members of the Mali lab for advice and help with experiments, and the Salk GT3 viral core for help with AAV production. This work was supported by UCSD Institutional Funds and NIH grants (R01HG009285, RO1CA222826, RO1GM123313). A.M. acknowledges a graduate fellowship from CONACYT and UCMEXUS. GFC thanks FAPESP (grant 2018/05778-3).

## AUTHOR CONTRIBUTIONS

A.M. conceived and performed experiments, analyzed data and wrote the manuscript. G.C. performed the BzATP experiments. F.A. performed experiments, analyzed data and wrote the manuscript. A.P., P.M., M.H. performed experiments. S.W. helped setup and design experiments. P.M. and T.Y. supervised the project, conceived and designed experiments, and wrote the manuscript. This article was prepared while Sarah A. Woller was employed at the University of California, San Diego. The opinions expressed in this article are the author’s own and do not reflect the view of the National Institutes of Health, the Department of Health and Human Services, or the United States Government.

## COMPETING INTERESTS

A.M., F.A., T.Y. and P.M. have filed patents based on this work. TY is a member of the SAB for Navega Therapeutics. A.M. is a co-founder of Navega Therapeutics. P.M. is a scientific co-founder of Shape Therapeutics, Boundless Biosciences, Seven Therapeutics, Navega Therapeutics, and Engine Biosciences. The terms of these arrangements have been reviewed and approved by the University of California, San Diego in accordance with its conflict of interest policies.

---

**Supplementary Figure 1:**
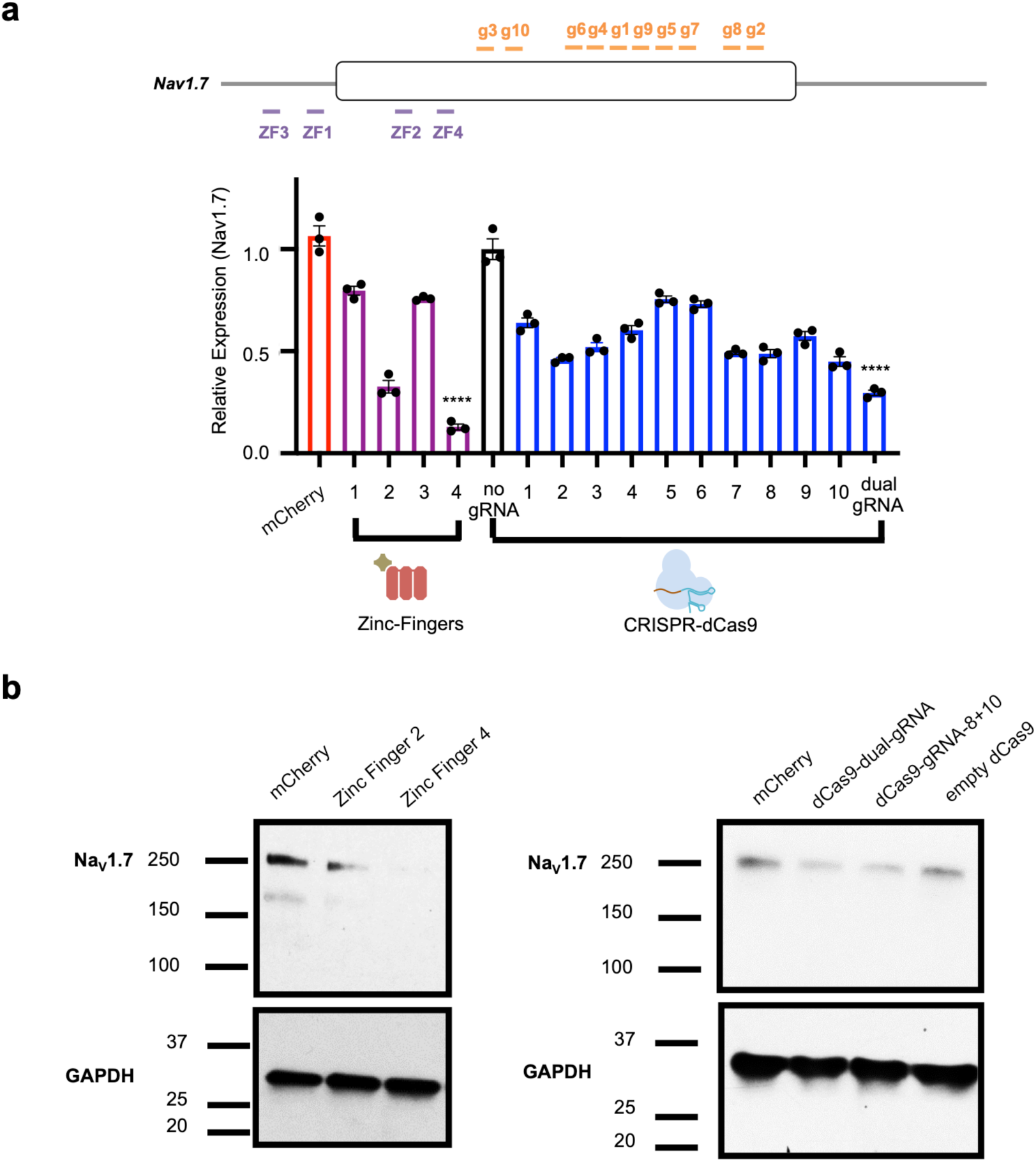
*In vitro* optimization of epigenetic genome engineering tools to enable Na_V_1.7 repression. **(a)** A panel of four zinc finger proteins and ten gRNAs were designed to target Na_V_1.7 in a mouse neuroblastoma cell line (Neuro2a) and were screened for repression efficacy by qPCR. A non-targeting gRNA (no gRNA) was used as a control for KRAB-dCas9 constructs targeting Na_V_1.7, while mCherry was used as a control for ZFP-KRAB constructs targeting Na_V_1.7 (n=3; error bars are SEM; one-way ANOVA; ****p < 0.0001). **(b)** *In vitro* western blotting of Na_V_1.7 in Neuro2a cells transfected with mCherry, Zinc-Finger-2-KRAB, Zinc-Finger-4-KRAB, KRAB-dCas9-no-gRNA, KRAB-dCas9-dual-gRNA (1+2), and KRAB-dCas9-gRNA-8+10.

**Supplementary Figure 2:**
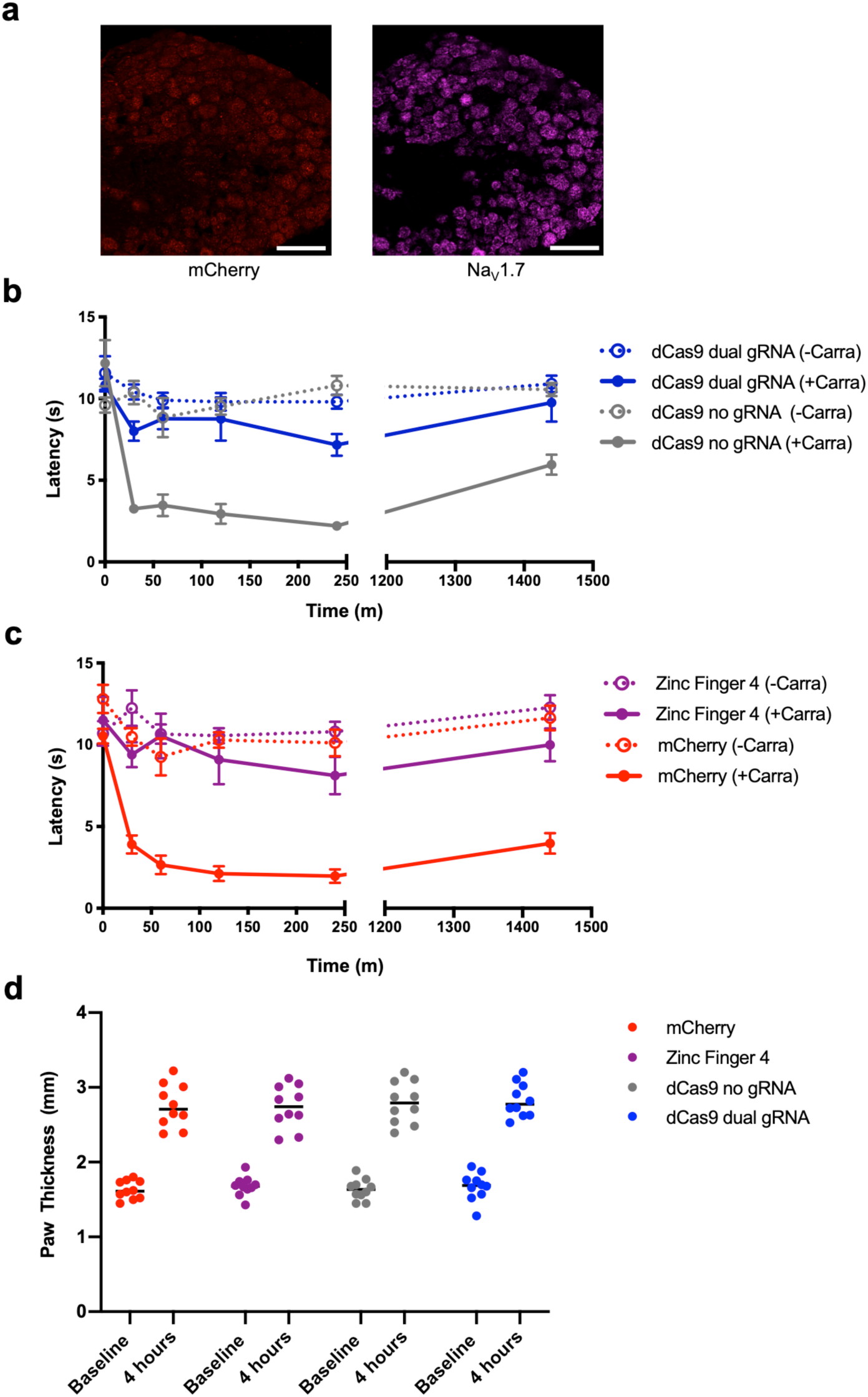
*In situ* repression of Na_V_1.7 leads to pain amelioration in a carrageenan model of inflammatory pain. **(a)** Confirming AAV9-mCherry transduction in mice DRG via RNA FISH (red= mCherry, pink= Na_V_1.7; scale bar=50µm). **(b)** Time course of thermal hyperalgesia after the injection of carrageenan (solid lines) or saline (dotted lines) into the hind paw of mice 21 days after i.t. injection with AAV9-KRAB-dCas9-no-gRNA and AAV9-KRAB-dCas9-dual-gRNA are plotted. Mean paw withdrawal latencies are shown. (n=10; error bars are SEM). **(c)** Time course of thermal hyperalgesia after the injection of carrageenan (solid lines) or saline (dotted lines) into the hind paw of mice 21 days after i.t. injection with AAV9-mCherry and AAV9-Zinc-Finger-4-KRAB are plotted. Mean paw withdrawal latencies are shown. (n=10; error bars are SEM). **(d)** Paw thickness of ipsilateral paws at baseline and four hours after carrageenan injection are plotted (n=10).

**Supplementary Figure 3:**
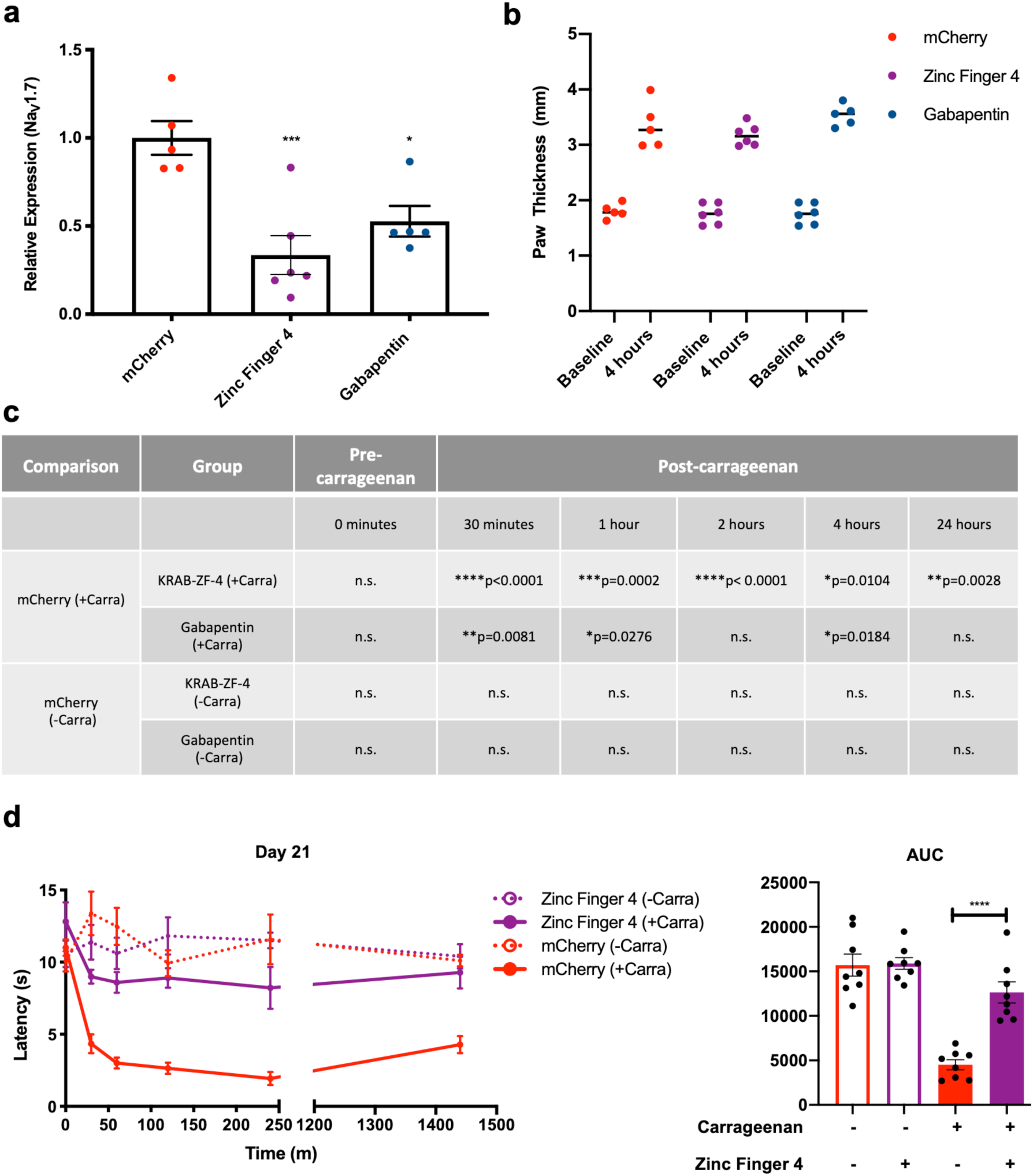
Evaluation of Zinc-Finger-KRAB in an inflammatory model of pain. **(a)** *In vivo* Na_V_1.7 repression efficiencies from treated mice DRG. Twenty-four hours after carrageenan administration, mice DRG (L4-L6) were harvested and Na_V_1.7 repression efficacy was determined by qPCR. (n=5 for mCherry and Gabapentin groups and n=6 for Zinc-Finger-4-KRAB group; error bars are SEM; one way ANOVA with Dunnet’s *post hoc* test; ***p = 0.0007, *p=0.0121). **(b)** Paw thickness of ipsilateral paws at baseline and four hours after carrageenan injection are plotted. **(c)** Significance of paw withdrawal latencies in mice receiving AAV9-Zinc-Finger-4-KRAB and gabapentin (100 mg/kg) as compared to AAV9-mCherry carrageenan-injected paw (negative control). Two-way ANOVA with Bonferroni *post hoc* test. **(d)** Independent repeat of experiment in **(a):** time course of thermal hyperalgesia after the injection of carrageenan (solid lines) or saline (dotted lines) into the hind paw of mice 21 days after i.t. injection with AAV9-mCherry and AAV9-Zinc-Finger-4-KRAB are plotted. Mean paw withdrawal latencies are shown. The AUC of the thermal-hyperalgesia time-course are plotted on the right panel. A significant increase in PWL is seen in the carrageenan-injected paws of mice injected with AAV9-Zinc-Finger-4-KRAB (n=8; error bars are SEM; Student’s t-test; ****p < 0.0001).

**Supplementary Table 1:**
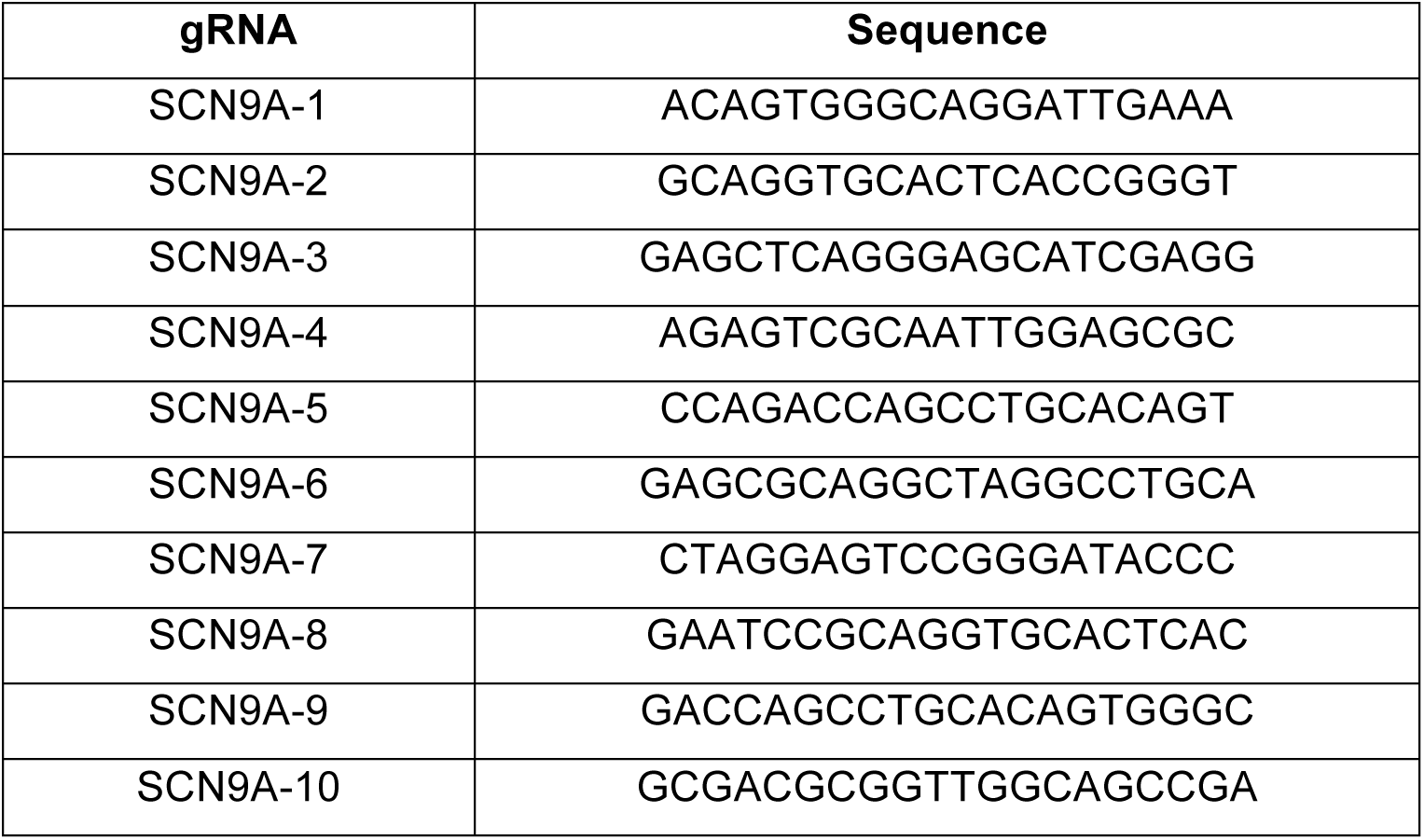
CRISPR-Cas9 guide RNA spacer sequences.

**Supplementary Table 2:**
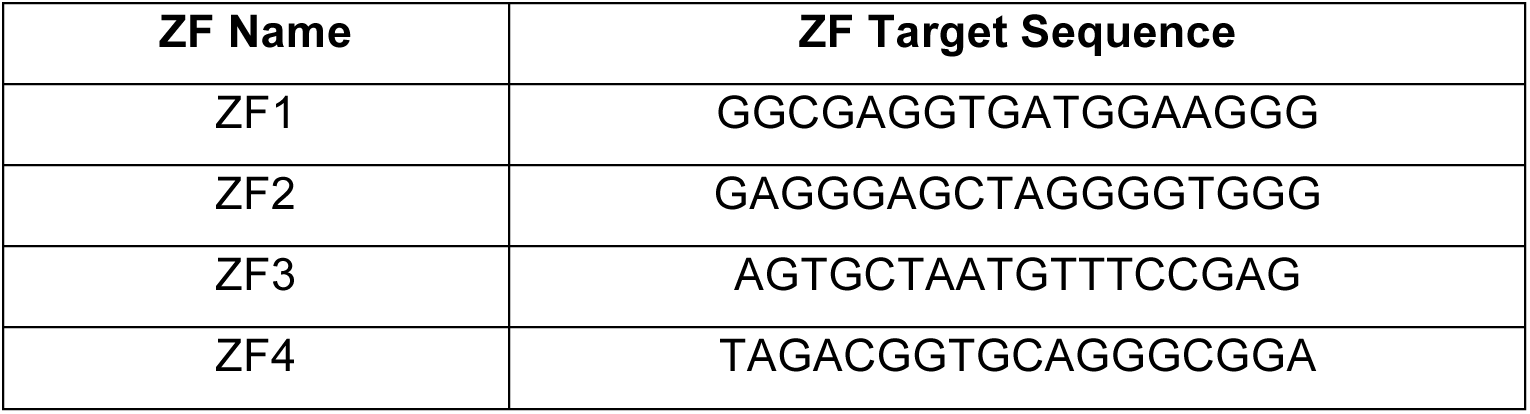
Zinc finger protein genomic target sequences.

## MATERIALS AND METHODS

### Vector Design and Construction

Cas9 and Zinc-Finger AAV vectors were constructed by sequential assembly of corresponding gene blocks (Integrated DNA Technologies) into a custom synthesized rAAV2 vector backbone. gRNA sequences were inserted into dNCas9 plasmids by cloning oligonucleotides (IDT) encoding spacers into AgeI cloning sites via Gibson assembly. gRNAs were designed utilizing an *in silico* tool to predict gRNAs^73^.

### Mammalian Cell Culture

Neuro2a cells were grown in EMEM supplemented with 10% fetal bovine serum (FBS) and 1% Antibiotic-Antimycotic (Thermo Fisher Scientific) in an incubator at 37°C and 5% CO_2_ atmosphere.

### Lipid-Mediated Cell Transfections

One day prior to transfection, Neuro2a cells were seeded in a 24-well plate at a cell density of 1 or 2E+5 cells per well. 0.5 µg of each plasmid was added to 25 µL of Opti-MEM medium, followed by addition of 25 µL of Opti-MEM containing 2 µL of Lipofectamine 2000. The mixture was incubated at room temperature for 15 min. The entire solution was then added to the cells in a 24-well plate and mixed by gently swirling the plate. Media was changed after 24 h, and the plate was incubated at 37°C for 72 h in a 5% CO_2_ incubator. Cells were harvested, spun down, and frozen at 80°C.

### Production of AAVs

Virus was prepared by the Gene Transfer, Targeting and Therapeutics (GT3) core at the Salk Institute of Biological Studies (La Jolla, CA) or in-house utilizing the GT3 core protocol. Briefly, AAV2/1, AAV2/5, and AAV2/9 virus particles were produced using HEK293T cells via the triple transfection method and purified via an iodixanol gradient. Confluency at transfection was between 80% and 90%. Media was replaced with pre-warmed media 2h before transfection. Each virus was produced in five 15 cm plates, where each plate was transfected with 10 µg of pXR-capsid (pXR-1, pXR-5, and pXR-9), 10 of µg recombinant transfer vector, and 10 µg of pHelper vector using polyethylenimine (PEI; 1 mg/mL linear PEI in DPBS [pH 4.5], using HCl) at a PEI:DNA mass ratio of 4:1. The mixture was incubated for 10 min at room temperature and then applied dropwise onto the media. The virus was harvested after 72 h and purified using an iodixanol density gradient ultracentrifugation method. The virus was then dialyzed with 1x PBS (pH 7.2) supplemented with 50 mM NaCl and 0.0001% of Pluronic F68 (Thermo Fisher Scientific) using 50-kDa filters (Millipore) to a final volume of ∼100 µL and quantified by qPCR using primers specific to the ITR region, against a standard (ATCC VR-1616): AAV-ITR-F: 5’ -CGGCCTCAGTGAGCGA-3’ and AAV-ITR-R: 5’ - GGAACCCCTAGTGATGGAGTT-3’

### Animals Experiments

All animal procedures were performed in accordance with protocols approved by the Institutional Animal Care and Use Committee (IACUC) of the University of California, San Diego. All mice were acquired from Jackson Laboratory. Two-month-old adult male C57BL/6 mice (25-30g) were housed with food and water provided *ad libitum*, under a 12 h light/dark cycle with up to 5 mice per cage. All behavioral tests were performed during the light cycle period.

### Intrathecal AAV Injections

Anesthesia was induced with 2.5% isoflurane delivered in equal parts O_2_ and room air in a closed chamber until a loss of the righting reflex was observed. The lower back of mice was shaven and swabbed with 70% ethanol. Mice were then intrathecally (i.t.) injected using a Hamilton syringe and 30G needle as previously described^104^ between vertebrae L4 and L5 with 5 µL of AAV for a total of 1E+12 vg/mouse. A tail flick was considered indicative of appropriate needle placement. Following injection, all mice resumed motor activity consistent with that observed prior to i.t. injection.

### Pain Models

#### Intraplantar carrageenan injection

Carrageenan-induced inflammation is a classic model of edema formation and hyperalgesia^105–107^. 21 days after AAV pre-treatment, anesthesia was induced as described above. Lambda carrageenan (Sigma Aldrich; 2% (W/V) dissolved in 0.9% (W/V) NaCl solution, 20 µL) was subcutaneously injected with a 30G needle into the plantar (ventral) surface of the ipsilateral paw. An equal amount of isotonic saline was injected into the contralateral paw. Paw thickness was measured with a caliper before and 4h after carrageenan/saline injections as an index of edema/inflammation. Hargreaves testing was performed before injection (t=0) and (t= 30, 60, 120, 240 minutes and 24 hours post-injection). The experimenter was blinded to the composition of treatment groups. Mice were euthanized after the 24-hour time point.

#### Paclitaxel-induced neuropathy

Paclitaxel (Tocris Biosciences, 1097) was dissolved in a mixture of 1:1:18 [1 volume ethanol/1 volume Cremophor EL (Millipore, 238470)/18 volumes of sterilized 0.9% (W/V) NaCl solution]. Paclitaxel injections (8 mg/kg) were administered intraperitoneally (i.p.) in a volume of 1 mL/100 g body weight every other day for a total of four injections to induce neuropathy (32 mg/kg), resulting in a cumulative human equivalent dose of 28.4–113.5 mg/m^2^ as previously described^67^. Behavioral tests were performed 24 hours after the last dosage.

#### Intrathecal BzATP injection

BzATP (2′(3′)-O-(4-Benzoylbenzoyl) adenosine 5′-triphosphate triethylammonium salt) was purchased from Millipore Sigma and, based on previous tests, was dissolved in saline (NaCl 0.9%) to final a concentration of 30 nmol. Saline solution was also used as a vehicle control and both were delivered in a 5 µL volume. Intrathecal injections were performed under isoflurane anesthesia (2.5%) by lumbar puncture with a 30-gauge needle attached to a Hamilton syringe.

### Behavioral tests

Mice were habituated to the behavior and to the experimental chambers for at least 30 min before testing. As a positive control, gabapentin (Sigma, G154) was dissolved in saline solution and injected i.p. at 100 mg/kg/mouse.

#### Thermal Withdrawal Latency (Hargreaves Test)

To determine the acute nociceptive thermal threshold, the Hargreaves’ test was conducted using a plantar test device (Ugo Basile, Italy)^108^. Animals were allowed to freely move within a transparent plastic enclosement (6 cm diameter × 16 cm height) on a glass floor 40 min before the test. A mobile radiant heat source was then placed under the glass floor and focused onto the hind paw. Paw withdrawal latencies were measured with a cutoff time of 30 seconds. An IR intensity of 40 was employed. The heat stimulation was repeated three times on each hind paw with a 10 min interval to obtain the mean latency of paw withdrawal. The experimenter was blinded to composition of treatment groups.

#### Tactile allodynia

For the BzATP pain model, tactile thresholds (allodynia) were assessed 30 minutes, 1, 2, 3, 6, 24 hours after the BzATP injection. For the Paclitaxel model, tactile thresholds (allodynia) were assessed 24 hours and 29 days after the last Paclitaxel injection. Forty-five minutes before testing, mice were placed in clear plastic wire mesh-bottom cages for acclimation. The 50% probability of withdrawal threshold was assessed using von Frey filaments (Seemes Weinstein von Frey anesthesiometer; Stoelting Co., Wood Dale, IL, USA) ranging from 2.44 to 4.31 (0.04–2.00 g) in an up-down method, as previously described^107^.

#### Cold allodynia

Cold allodynia was measured by applying drops of acetone to the plantar surface of the hind paw as previously described^109,110^. Mice were placed in individual plastic cages on an elevated platform and were habituated for at least 30 min until exploratory behaviors ceased. Acetone was loaded into a one mL syringe barrel with no needle tip. One drop of acetone (approximately 20 µL) was then applied through the mesh platform onto the plantar surface of the hind paw. Care was taken to gently apply the bubble of acetone to the skin on the paw without inducing mechanical stimulation through contact of the syringe barrel with the paw. Paw withdrawal time in a 60s observation period after acetone application was recorded. Paw withdrawal behavior was associated with secondary animal responses, such as rapid flicking of the paw, chattering, biting, and/or licking of the paw. Testing order was alternated between paws (i.e. right and left) until five measurements were taken for each paw. An interstimulation interval of 5 minutes was allowed between testing of right and left paws.

### Tissue collection

After the 24-hour time carrageenan time point, spinal cords were collected via hydroextrusion (injection of 2 mL of iced saline though a short blunt 20 gauge needle placed into the spinal canal following decapitation). After spinal cord tissue harvest, the L4-L6 DRG on each side were combined and frozen as for the spinal cord. Samples were placed in DNase/RNase-free 1.5 mL centrifuge tubes, quickly frozen on dry ice, and then stored at 80°C for future analysis.

### Gene Expression Analysis and qPCR

RNA from Neuro2a cells was extracted using RNeasy Kit (QIAGEN; 74104) and from DRG using RNeasy Micro Kit (QIAGEN; 74004). cDNA was synthesized from RNA using Protoscript II Reverse Transcriptase Kit (NEB; E6560L). Real-time PCR (qPCR) reactions were performed using the KAPA SYBR Fast qPCR Kit (Kapa Biosystems; KK4601), with gene-specific primers in technical triplicates and in biological triplicates (Neuro2a cells). Relative mRNA expression was normalized to GAPDH levels and fold change was calculated using the comparative CT (ΔΔCT) method and normalized to GAPDH. Mean fold change and SD were calculated using Microsoft Excel.

### Western Blot

Neuro2a cells were thawed and protein extraction was performed with RIPA buffer (25mM Tris•HCl pH 7.6, 150mM NaCl, 1% NP-40, 1% sodium deoxycholate, 0.1% SDS; Thermo Fisher 89900) supplemented with protease inhibitors (Sigma P8849). Total protein was quantified with BCA protein assay kit (Thermo Fisher 23225), and 40 µg of protein were loaded into 4-15% polyacrylamide gels (BioRad 4561085). Proteins were transfer to a PVDF membrane (Thermo Fisher IB401001) and the membrane was blocked with 5% (W/V) blotting-grade blocker (Biorad 1706404) dissolved in TBS-T (Thermo Fisher, 28358 supplemented with 0.1% (V/V) Tween-20; BioRad 1610781). Membranes were then incubated overnight at 4°C with primary antibodies: anti-Na_V_1.7 diluted 1:1000 (Abcam; ab85015) and anti-GAPDH (Cell Signaling, 2118) diluted 1:4000. Membranes were then washed three times with TBS-T and incubated for 1 h at room temperature with anti-rabbit horseradish-peroxidase-conjugated secondary antibody (Cell Signaling, 7074) diluted 1:20000. After being washed with TBST, blots were visualized with SuperSignal West Femto Chemiluminescent Substrate (Thermo Fisher) and visualized on an X-ray film.

### RNAscope ISH Assays

The mCherry, and Na_V_1.7 probes were designed by Advanced Cell Diagnostics (Hayward, CA). The mCherry probe (ACD Cat# 404491) was designed to detect 1480– 2138 bp (<u>KF450807.1</u>, C1 channel), and the Na_V_1.7 (ACD Cat#313341), was designed to detect 3404–4576 bp of the *Mus musculus* Na_V_1.7 mRNA sequence (<u>NM_018852.2</u>, C3 channel). Before sectioning, DRG were placed into 4% PFA for 2 hours at room temperature, followed by incubation in 30% sucrose overnight at 4°C. Tissues were sectioned (12 µm thick) and mounted on positively charged microscopic glass slides (Fisher Scientific). All hybridization and amplification steps were performed following the ACD RNAscope V2 fixed tissue protocol. Stained slides were coverslipped with fluorescent mounting medium (ProLong Gold Anti-fade Reagent P36930; Life Technologies) and scanned into digital images with a Zeiss 880 Airyscan Confocal at 20x magnification. Data was processed using ZEN software (manufacturer-provided software).

### Statistical analysis

Results are expressed as mean +/− standard error (SE). Statistical analysis was performed using GraphPad Prism (version 8.0, GraphPad Software, San Diego, CA, USA). Results were analyzed using Student’s t-test (for differences between two groups), one-way ANOVA (for multiple groups), or two-way ANOVA with the Bonferroni *post hoc* test (for multiple groups time-course experiments). Differences between groups with p < 0.05 were considered statistically significant.

